# Distinct response selectivity changes in primary visual and parietal cortex during visual discrimination learning

**DOI:** 10.1101/2025.10.24.684194

**Authors:** John P. McClure, Mátyás Váradi, Jasper Poort

## Abstract

When animals learn the behavioral relevance of sensory features, response selectivity in primary sensory areas increases for those features. However, the effect of learning on neuronal activity in higher-level areas associated with decision-making in the parietal cortex, compared to primary sensory cortex, is not yet known. We used two-photon calcium imaging to determine how learning modifies neural representations in primary visual cortex (V1) and posterior parietal cortex (PPC) in a visual go/no-go orientation discrimination task. We found that behavior improvements after learning were associated with increased neuronal selectivity in both V1 and PPC. The increased selectivity in PPC was mainly driven by neurons preferring the rewarded go stimulus, while, in V1, neurons increased their selectivity for both the rewarded go and unrewarded no-go stimulus. Furthermore, feature preference was robust in V1 neurons but reorganized in PPC after learning. Finally, feature preference was preserved across contexts in V1 but not in PPC after learning, where many neurons switched their preference to the rewarded feature during active task engagement. Our results demonstrate that learning a visually-guided discrimination task increased information about relevant sensory features through distinct changes in the bottom and top levels of the visual cortical hierarchy. Visual cortex neurons encode visual features with increased reliability but with preserved feature preferences, while parietal neurons reorganize their feature preferences in a task-specific manner.

**Highlights:**

- V1 and PPC response selectivity increases after visual discrimination learning.
- Selectivity increase in PPC for rewarded features, but in V1 for both rewarded and unrewarded features.
- Feature preference robust in V1 but altered in PPC across learning.
- Feature preference after learning preserved across contexts in V1 but not PPC.

## Introduction

Animals have limited brain capacity and cannot process all sensory features in their environment. Learning enables selection and prioritized processing of features relevant for decision-making and action in a hierarchy of sensory brain regions. However, the effect of learning on neuronal feature representations in low-level sensory cortex compared to higher-level areas associated with decision-making is not understood. We therefore investigated the effects of visual discrimination learning in mice on the primary visual cortex and posterior parietal cortex – two regions at opposite ends of the cortical hierarchy.

The primary visual cortex (V1) contains precise representations of visual features, including stimulus orientation (Hubel and Wiesel, 1962; Niell and Stryker, 2008). V1 projects onto higher-level visual and association areas for higher-order visual processing during visually-guided decision-making tasks (Felleman and Essen, 1991; Wang and Burkhalter, 2007; Zhang et al., 2016). Association areas such as the posterior parietal cortex (PPC) are thought to play a key role in translating visual and other sensory representations into decisions and motor output (Colby and Goldberg, 1999; Andersen and Buneo, 2002; Hovde et al., 2019; Lyamzin and Benucci, 2019).

Previous studies found that both V1 and PPC are involved in visual discriminations: optogenetic silencing of V1 (Glickfeld et al., 2013; Poort et al., 2015) and PPC (Goard et al., 2016) disrupts visual discriminations. At the neuronal level, neurons in V1 increase their response selectivity for task-relevant features during visual discrimination learning when mice learn to respond to a rewarded visual stimulus and ignore a non-rewarded visual stimulus (Poort et al., 2015; Jurjut et al., 2017; Henschke et al., 2020; Goltstein et al., 2013). In PPC neurons in trained animals, activity is biased to task-relevant rewarded features (Goard et al., 2016; Pho et al., 2018).

In this study, we addressed the following key unresolved questions. First, what is the effect of learning on feature representations in PPC compared to V1? Second, how do learning effects in both regions depend on existing feature preferences of cells? Third, how specific are learning effects to the task that the animal performs, and what is the difference between task-relevant rewarded and unrewarded features?

To address these questions, we used two-photon calcium imaging to measure activity of layer 2/3 neuronal populations in V1 and PPC while mice learned to discriminate between two visual orientations, with one orientation (go stimulus) being rewarded. We found that both PPC and V1 neurons displayed increased selectivity after learning. However, the increase in PPC selectivity was primarily driven by go-preferring neurons, while in V1, both go- and no-go-preferring neurons showed increased selectivity. The feature preference of V1 neurons was robust across learning and task context, while it was highly dynamic in PPC with neurons switching their preference to the rewarded features compared to their pre-learning preferences and compared to their responses outside of the task context. Our results reveal that learning a visually-guided discrimination task enhances information about relevant sensory features through distinct patterns of changes in the bottom and top levels of the visual cortical hierarchy, with an enhanced but robust representation of task-relevant features and a winner-takes-all saliency representation in the parietal cortex biased to rewarded features.

## Results

To investigate neural changes in V1 and PPC during visual discrimination learning, we trained mice on a go/no-go visual discrimination task (Figure 1A). Head-fixed mice were positioned on a cylindrical treadmill. Trials were initiated by the animals’ running, and they were rewarded for licking in response to the go stimulus (static 45° or -45° grating) while no reward was delivered for licking during presentation of the no-go stimulus (with orientation orthogonal to the go-stimulus). Task performance was quantified by calculating the behavioral d-prime from the hit and false alarm rate (see STAR Methods). Mice learned on average within 3.5 weeks to lick in response to the presentation of the rewarded grating and withhold licking for the non-rewarded grating (Figure 1C, d-prime in the last session > 1.75 corresponding to on average 85% accuracy post-learning, average behavioral d-prime pre-learning -0.29, post learning 2.61, Wilcoxon signed-rank test, p = 4.3 × 10^-4^, N=16).

**Figure 1.**
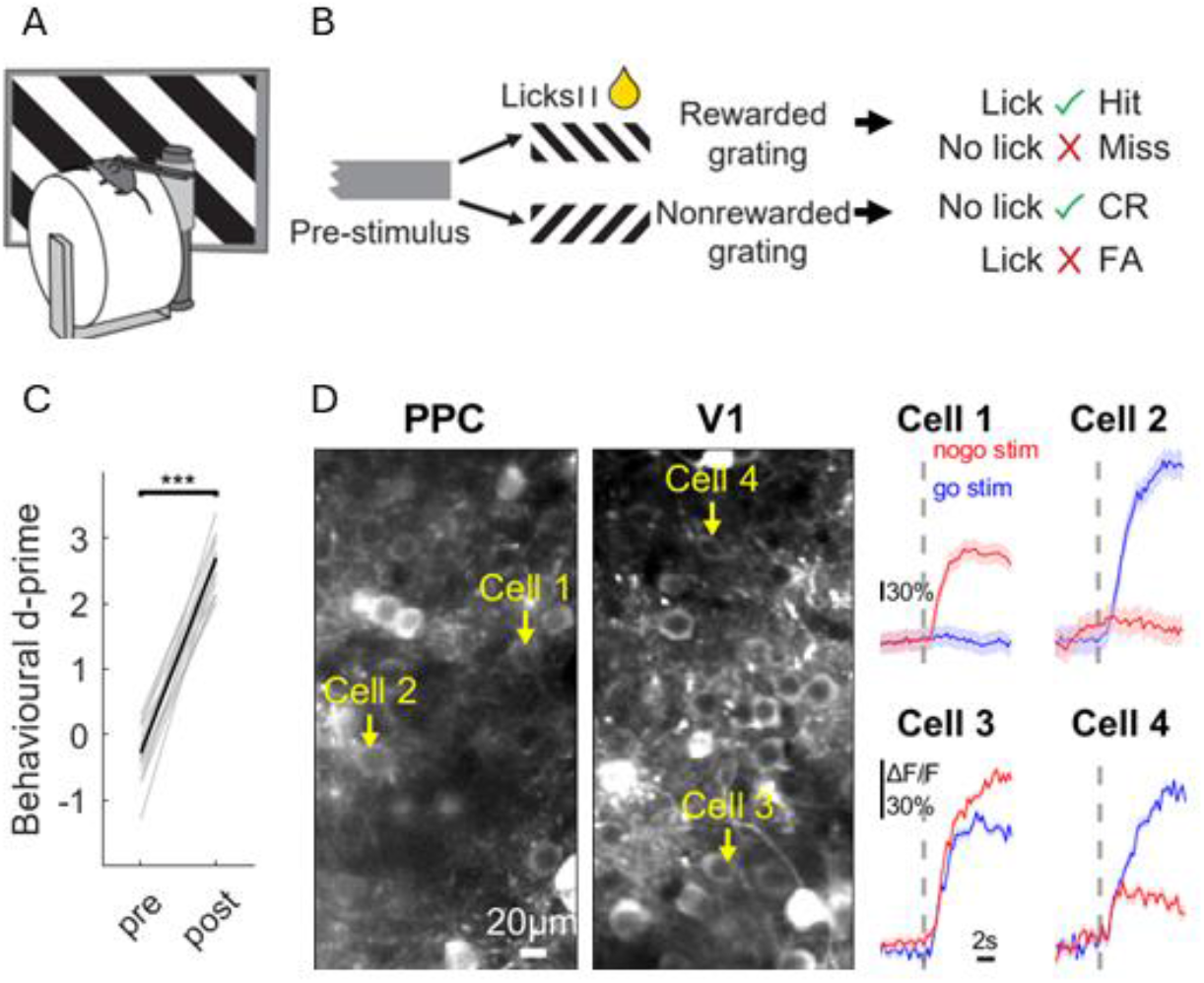
Learning of visual discrimination task and response selectivity for visual stimuli in PPC and V1. **A)** Schematic of the go/no-go task setup (Kukovska and Poort, 2024). Mice were head-fixed on cylindrical treadmill. Stimuli were presented on monitor to the right eye (contralateral to the imaged V1 or PPC). Licking spout in front of mouse delivered strawberry milk reward. **B)** Schematic of visual discrimination task. Mice initiated presentation of go or no-go stimulus by crossing running speed threshold during pre-stimulus baseline period with grey background (>0.5 s). Go (45 or -45 degrees orientation, counterbalanced across animals) or no-go static visual grating (with orientation orthogonal to go stimulus) was presented for 2 s. Licks were detected during response window (1-2 s post stimulus onset) to determine whether a trial was a hit, miss, false alarm (FA) or correct rejection (CR). **C)** Pre-and post-learning task performance. Solid line, d’ averaged across mice. Grey lines, individual mice (***p<0.001, Wilcoxon signed-rank test, N = 16). **D)** Example neurons from PPC and V1. Left, PPC and V1 two-photon imaging planes. Right, average neuronal responses to the go (blue trace) and no-go (red trace) stimulus during discrimination task. Shading, SEM. Dashed lines indicate stimulus onset.

### V1 and PPC selectivity changes during learning

Before animal training, we expressed calcium indicator GCaMP7 (Dana et al., 2019) in V1 and PPC using viral vectors. We measured responses in layer 2/3 V1 and PPC neurons (Figure 1D) using two-photon calcium imaging while animals learned and performed the visual discrimination task.

Neurons from each area showed different degrees of responses to the go and no-go stimulus. After learning, both V1 and PPC neurons showed an increased response difference for the go and no-go stimulus (Figure 2A). To quantify this, we calculated the stimulus selectivity (Figure 2B, difference in the mean responses to the two grating stimuli, normalized by the response variability; positive values indicate preference for the go stimulus and negative values indicate preference for the no-go stimulus, see STAR Methods). The absolute selectivity quantifies how well neurons discriminate irrespective of their preference. Neurons in both V1 and PPC showed increased absolute stimulus selectivity after learning (Figure 2C, V1, absolute selectivity pre-learning 0.28 ± 0.39, post-learning 0.35 ± 0.39, mean ± SD, N_pre_=2608 cells N_post_=2130, p < 0.0001, bootstrap test; PPC pre 0.25 ± 0.33, post 0.29 ± 0.37, N_pre_=1726 cells, N_post_=1726, p = 0.003; see also Supplementary Figure 1A). The increase in V1 was driven by both go- and no-go preferring cells (Figure 2D, go-preferring cells, absolute selectivity pre-learning 0.28 ± 0.38, post-learning 0.31 ± 0.31, mean ± SD, N_pre_=1400, N_post_=1156, p = 0.0002, bootstrap test; no-go preferring cells absolute selectivity pre 0.28 ± 0.40, post 0.39 ± 0.45, N_pre_=1207, N_post_=974, p < 0.0001). In contrast, the increase in PPC was driven by the go-preferring cells (Figure 2D, go cells absolute selectivity pre 0.25 ± 0.31, post 0.32 ± 0.34, mean ± SD, N_pre_=368, N_post_=1106, p = 0.0002, bootstrap test; no-go cells absolute selectivity pre 0.25 ± 0.36, post 0.24 ± 0.40, p = 0.35, N_pre_=326, N_post_=620)

**Figure 2.**
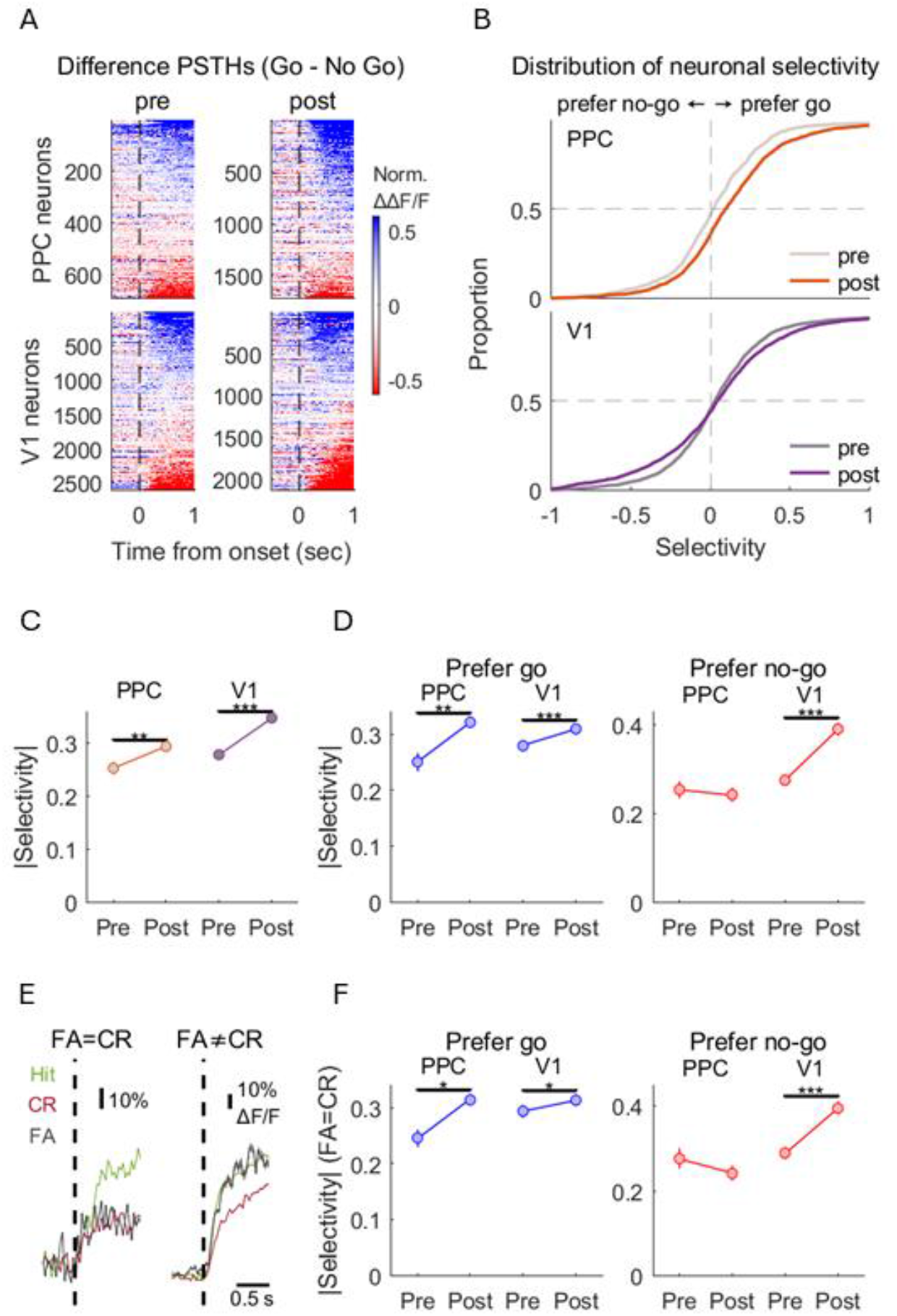
PPC neurons become more selective for rewarded go stimulus, while V1 neurons become more selective for both go and no-go stimuli after visual discrimination learning. **A)** Response difference between the rewarded (go) and non-rewarded (no-go) stimulus divided by baseline variance (-1 to 0 from stimulus onset) before (left column) and after learning (right column) in PPC (top row) and V1 (bottom row). Cells sorted by average amplitude 0–1 s from stimulus onset. **B)** Cumulative distribution of neuronal selectivity before (faint line) and after learning (dark line) in PPC (top panel) and V1 (bottom panel). Positive values: cells prefer rewarded (go) grating; negative values: cells prefer non-rewarded (no-go) grating. **C)** Average absolute selectivity before and after learning in PPC and V1. **D)** Average absolute selectivity before and after learning in PPC and V1 for cells preferring the rewarded go (selectivity index > 0, left panel) and non-rewarded no-go stimulus (selectivity index < 0, right panel). **E)** Average grating responses of two example cells classified as behaviorally not modulated (left; responses are similar in FA and CR trials, p ≥ 0.05, Wilcoxon rank-sum test) and behaviorally modulated (right; responses are different in FA and CR trials, p < 0.05, Wilcoxon rank-sum test). Shading, SEM. **F)** Average absolute selectivity before and after learning in PPC and V1 for cells not modulated by behavior (responses are similar in FA and CR trials, p ≥ 0.05, Wilcoxon rank-sum test; minimum three FA and three CR trials). Left panel, go-preferring cells (selectivity index > 0), right panel, no-go-preferring cells (selectivity index < 0). **C, D**, and **F)** Error bars, SEM. *p<0.05, **p<0.01, ***p<0.001, bootstrap test (see main text for details).

To determine whether the increased selectivity in PPC and V1 after learning was due to differences in behavior, we performed a control analysis comparing activity during correct rejection trials (no lick) and false alarm trials (erroneous licks). First, we determined for each cell whether the stimulus response was the same during correct rejections and false alarms or not (Figure 2E). Next, we repeated our analysis after only including cells whose responses to the stimuli were the same irrespective of the behavior (Figure 2F). The results were the same as before: V1 showed increased selectivity for both go and no-go preferring cells (go-preferring cells, absolute selectivity pre-learning 0.29 ± 0.39, post-learning 0.31 ± 0.33, mean ± SD, N_pre_= 1173, N_post_= 923, p = 0.03, bootstrap test; no-go preferring cells absolute selectivity pre 0.29 ± 0.43, post 0.40 ± 0.44, N_pre_= 938, N_post_= 775, p < 0.0001), while PPC only showed in increased response selectivity for the rewarded go-stimulus (go-preferring cells, absolute selectivity pre-learning 0.25 ± 0.27, post-learning 0.31 ± 0.34, mean ± SD, N_pre_= 304, N_post_= 900, p = 0.02, bootstrap test; no-go preferring cells absolute selectivity pre 0.28 ± 0.37, post 0.24 ± 0.42, N_pre_= 246, N_post_= 548, p = 0.06). Furthermore, results were also similar when only including cells with similar responses during hit and miss trials (Supplementary Figure 1B). As an additional control we used a generalized linear model (GLM) to identify cells modulated by licking and running. After excluding 50% of cells most modulated by these predictors, results were similar (Supplementary Figure 1C-D), indicating that the neural selectivity changes in V1 and PPC were not due to changes in licking or running. Interestingly, GLMs including visual, licking and running predictors revealed that while V1 was more modulated by stimulus presentation, PPC neurons were multiplexing sensory information and motor-output (Supplementary Figure 2A-B).

### Stimulus selectivity stable in V1 but changes in PPC across learning

To understand the nature of the selectivity changes in V1 and PPC, we compared the selectivity before and after learning in the same cells tracked across learning in V1 and PPC (Figure 3A-B). The selectivity before and after learning was relatively stable in V1 (Figure 3C, Pearson R=0.43, p = 3 × 10^-20^), indicating that V1 neurons preserve their feature preference across learning. However, in contrast, the correspondence between the pre- and post-learning selectivity in PPC was strikingly low (R=0.05, p = 0.32), indicating that PPC neurons can override their initial stimulus preference when they become biased to the rewarded go-stimulus after learning.

**Figure 3.**
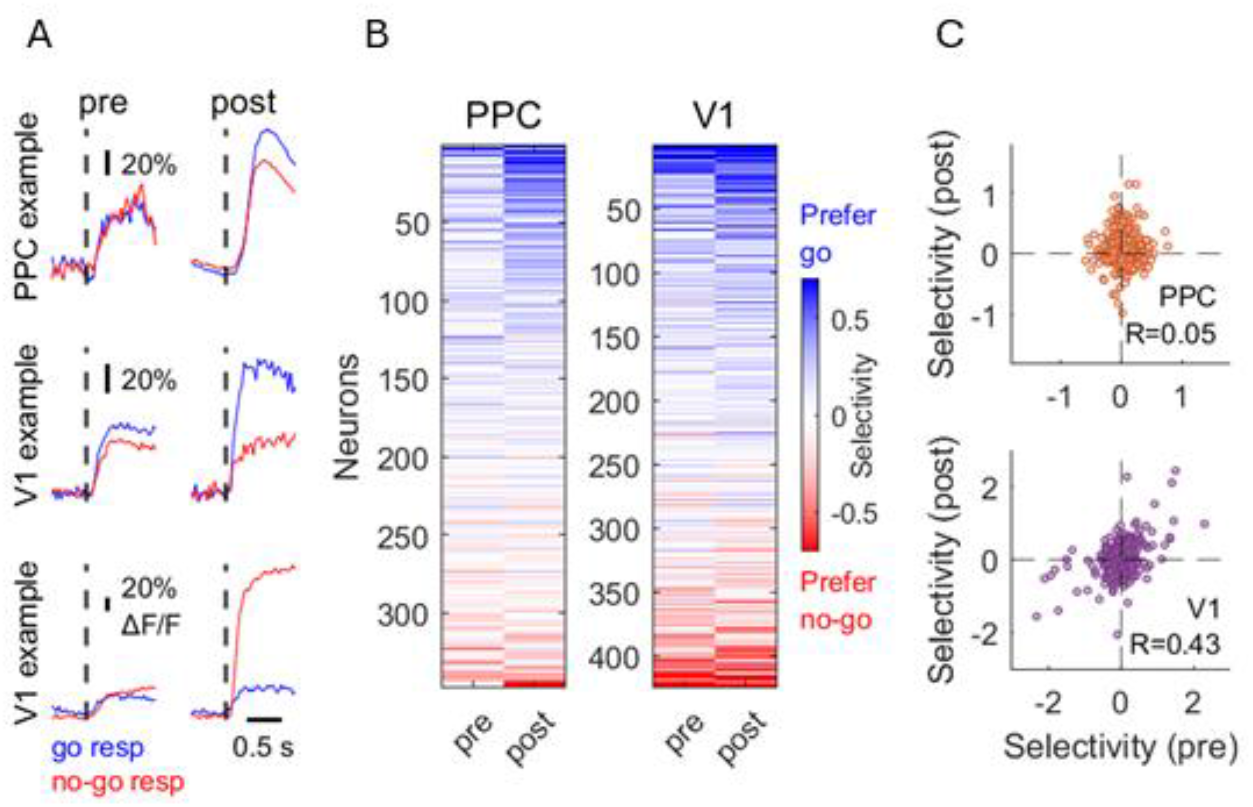
Stimulus preference remains stable in V1 but changes in PPC during learning. **A)** Average grating responses of three example neurons (from PPC: top; from V1: middle and bottom) before (left column) and after learning (right column). **B)** Stimulus selectivity of the same neurons (rows) before and after learning (columns) in PPC (left) and V1 (right). Cells were ordered by their mean pre- and post-learning selectivity (PPC: n = 10 mice; V1: n = 9 mice). **C)** Relationship between pre- and post-learning neuronal selectivity in PPC (top; p = 0.32) and V1 (bottom; p = 3 × 10^-20^). R: Pearson correlation coefficient.

### Stronger task-dependence of selectivity in PPC than in V1

In addition to measuring activity in V1 and PPC during active task performance, we also measured orientation tuning during passive viewing. During orientation tuning, we presented the go and no-go stimuli and an additional set of task-irrelevant orientations (Figure 4A). To further understand the changes in stimulus selectivity across learning, we compared the selectivity for the task-relevant go and no-go stimuli during two contextual conditions: during the task and outside of the task (during passive viewing). Before learning, both V1 and PPC showed stable stimulus selectivity in these two contexts (Figure 4B, left, pre learning V1 R=0.55, PPC: R=0.48, both p < 1 × 10^-15^). Although learning reduced the correspondence between the selectivity in the two contexts in V1 (Figure 4B, right bottom, R=0.38, p < 1 × 10^-15^), the correlation remained relatively high after learning. However, strikingly, the correspondence between the two contexts was greatly reduced after learning in PPC (Figure 4B, right top, R=0.09, p = 2.22 × 10^-6^). The results were the same after only including neurons with similar activity in correct rejection and false alarm, or in Hit and Miss trials (Supplementary Figure 3A-B), or after excluding the top half of the cells with the highest licking- or running-related GLM variance (Supplementary Figure 3C-D). Thus, in summary, V1 neurons display relatively robust feature preference while PPC neurons flexibly adapt their feature preferences after learning and during task performance.

**Figure 4.**
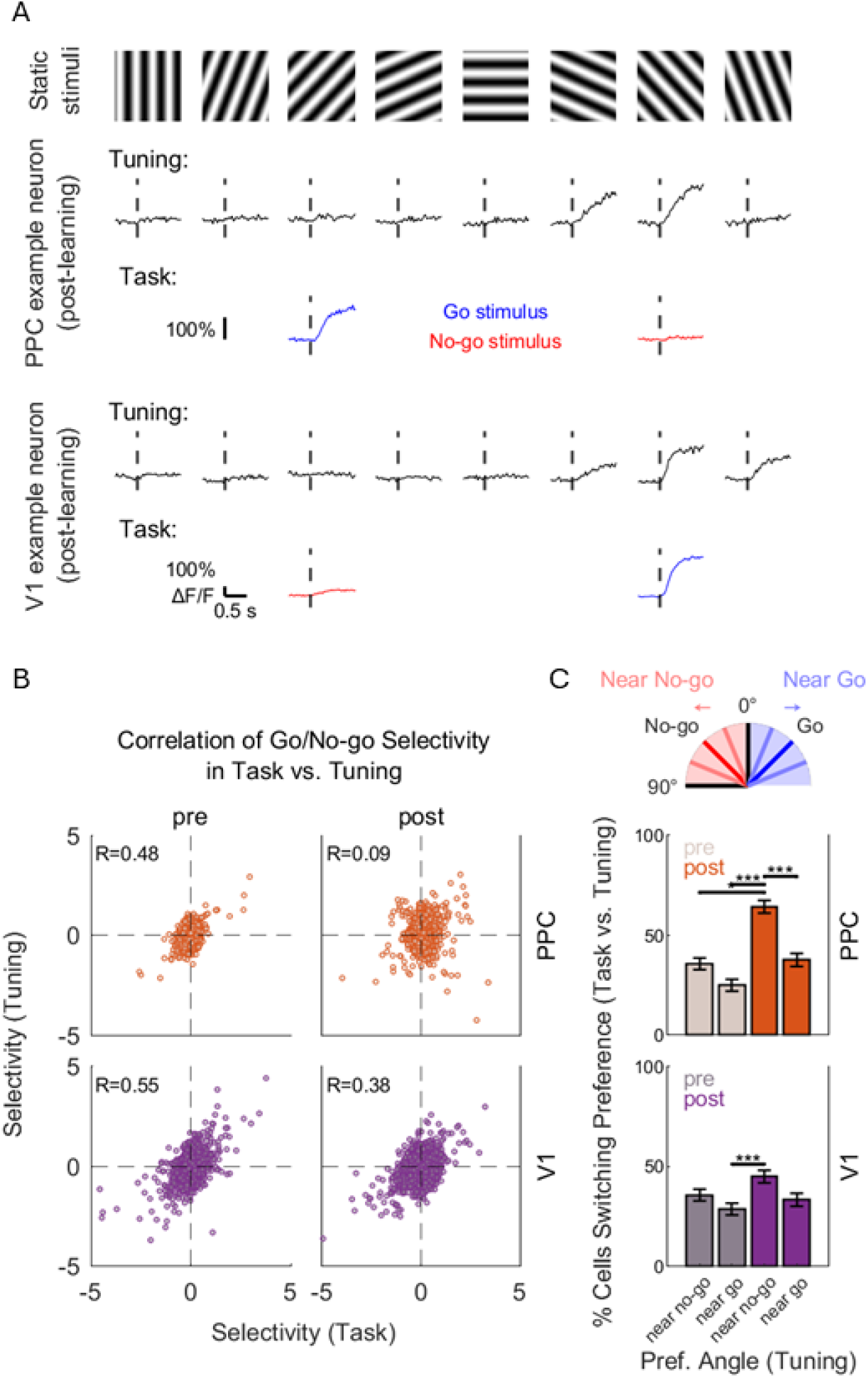
After learning, PPC neurons are biased towards the rewarded stimulus in a context dependent manner. **A)** Average response of an example neuron from PPC (middle row) and V1 (bottom row) to the static grating stimuli (top row) in orientation tuning (black traces) and go/no-go task (coloured traces). Blue and red colours indicate which stimulus was go and no-go in the discrimination task, respectively. Shading, SEM. **B)** Correlation of neuronal selectivity between the go and the no-go stimulus in orientation tuning and go/no-go task before (left column) and after learning (right column) in PPC (top row) and V1 (bottom row). R: Pearson’s correlation coefficient. PPC, pre-learning: N = 1353 cells, p = 5.87 × 10^-79^; PPC, post-learning: N = 2492 cells, p = 2.22 × 10^-6^; V1, pre-learning: N = 2718 cells, p = 1.95 × 10^-210^; V1, post-learning: N=2467, p=1.62 × 10^-85^. **C)** Proportions of cells switching orientation preference from tuning to task across different orientation preferences. Orientation preferences in tuning were grouped into ‘near go’ and ‘near no-go’ based on their proximity to task-relevant stimuli (top schematic; cells preferring 0° or 90° were excluded). A switching cell was defined as a neuron with ‘near no-go’ preference in tuning turning go-preferring (selectivity > 0) in task, or vice versa. pre, faint; post, dark. * p < 0.05, *** p < 0.001, bootstrap test. Error bars, bootstrap standard error (see Methods).

To further understand this change, we determined how cells modified their orientation preference from passive viewing to task. We determined the preferred orientations of cells, and grouped them into ‘near go’ and ‘near no-go’ neurons, based on which task-relevant grating their preferred angles were closer to in orientation tuning. Then, in each group, we calculated the proportion of those cells which switched orientation preference in the go/no-go task: a cell was defined as ‘switching’ if it preferred ‘near no-go’ orientations in passive viewing, but preferred go (selectivity > 0) in task, or vice versa (Figure 4C; PPC pre, near no-go: 36% of N=28; PPC, pre, near go: 25% of N=40; PPC, post, near no-go: 64% of N=112; PPC, post, near go: 38% of N=223; V1, pre, near no-go: 36% of N=266; V1, pre, near go: 29% of N=234; V1, post, near no-go: 45% of N=247; V1, post, near go: 33% of N=210). Strikingly, after learning, switching cells were significantly more common (over 50%) among ‘near no-go’ cells in PPC compared to pre-learning or ‘near go’ cells. Thus, in trained mice, most PPC neurons with a ‘near no-go’ preference became go-preferring in the visual discrimination task. In V1, switching cells were also more common in the ‘near no-go’ group in post-learning, but their proportion was only significantly different from the ‘near go’ neurons in pre-learning. Thus, visually-guided discrimination learning leads to a task-dependent reorganization of orientation preference in PPC.

## Discussion

In this study, we investigated the effect of learning on feature representations in two regions at opposite ends of the visual cortical hierarchy. We measured neuronal activity before and after learning and within different contexts. Therefore, we were able to link response changes to the original feature preferences of neurons and determine the effect of task context. We found that learning increases response selectivity of neurons in both the primary visual cortex (V1) and posterior parietal cortex (PPC). While the increase in V1 was seen for both go-preferring and no-go preferring cells, it was only observed in PPC cells preferring the rewarded go-stimulus. Feature preference was robust in V1 across learning, but it was reorganized in PPC after learning with distinct feature preference emerging during task performance.

V1 is located at the bottom of visual cortical hierarchy, as the first recipient of inputs from the dorsal lateral geniculate nucleus and projecting to a multitude of higher-level visual areas (Felleman and Essen, 1991; Wang and Burkhalter, 2007; Glickfeld and Olsen, 2017; Seabrook et al., 2017). In return, V1 receives a wide-range of top-down inputs, including projections form parietal cortex (Zhang et al., 2016; Morimoto et al., 2021). At the behavioral level, optogenetic inactivation of primary visual cortex disrupts basic visual detection and discrimination (Glickfeld et al., 2013; Poort et al., 2015; Goard et al., 2016). Cells in primary visual cortex have precise feature tuning, for example for stimulus orientation. Previous studies demonstrated that response selectivity of cells in V1 is enhanced after learning, and that V1 selectivity is correlated with behavioral performance (Poort et al., 2015). Here we replicated these findings and found an increase in response selectivity for both the rewarded go-stimulus and the non-rewarded no-go stimulus. In addition, while we see an enhancement with learning, we show that V1 feature preferences are relatively robust. Furthermore, our results demonstrate strong correlation of feature preferences within and outside of the task. These findings demonstrate that while primary visual cortex activity is modified with learning, it continues to provide a faithful representation of stimulus features across different contexts. This simultaneously allows for veridical encoding of stimulus features and stimulus saliency (Roelfsema, 2006). We found that V1 neurons were most strongly modulated by visual stimuli. This corroborates previous studies showing that V1 responses, in the presence of visual cues, primarily rely on visual information compared to other signals including motor variables or task context (Pakan et al., 2018; Pho et al., 2018). At the same time, we also identified behaviorally modulated V1 neurons, consistent with previous studies demonstrating, for example, modulation by locomotion (Niell and Stryker, 2010; Saleem et al., 2013; Pakan et al., 2016, 2018).

PPC sits at a relatively high level of the visual cortical hierarchy, and integrates inputs from early visual areas including V1, other sensory modalities, other association cortex including prefrontal cortex, and motor regions (Harvey et al., 2012; Zhang et al., 2016; Hovde et al., 2019; Lyamzin and Benucci, 2019). Optogenetic silencing of PPC disrupts performance of a memory-guided decision-making or visual discrimination task (Harvey et al., 2012; Goard et al., 2016; Licata et al., 2017). Additionally, previous studies have implicated PPC in visually-guided navigation (McNaughton et al., 1994; Krumin et al., 2018; Sorrell et al., 2025; Whitlock, 2017). To enable direct comparison with previous studies on mouse PPC, we used the same stereotaxic coordinates, which correspond to the rostro-medial tip of visual area A (Lyamzin and Benucci, 2019). Similar to previous studies, we found that PPC neurons compared to V1 show stronger multiplexing of visual- and motor-related variables such as running and licking (Pho et al., 2018), supporting the role of PPC in sensorimotor transformation (Colby and Goldberg, 1999; Andersen and Buneo, 2002; Hovde et al., 2019; Lyamzin and Benucci, 2019). PPC also showed distinct learning patterns from V1, as the selectivity increase was primarily driven by go-preferring neurons. By matching cells across learning, we demonstrate that PPC cells that originally prefer other orientations profoundly change their preference in a way that is not observed in V1. These results indicate that PPC activity is much more flexible than in V1.

Interestingly, before learning, we found that visual response selectivity was highly correlated during the task and when animals passively viewed the same features. However, response selectivity only increased for the rewarded features after learning. Furthermore, after learning, response selectivity during the task was not correlated anymore with selectivity observed during passive viewing. These results match with previous studies highlighting distinct responses in PPC during two distinct tasks (Lee et al., 2022), and previous studies showing a bias for task-relevant rewarded features in trained animals (Goard et al., 2016; Pho et al., 2018). These results are also consistent with the parietal cortex containing a saliency map with a winner-take-all mechanism (Itti and Koch, 2001).

In summary, our results highlight a fundamental difference in learning-related changes in early sensory cortex compared to higher-level association cortex. Learning enhances neuronal selectivity in primary sensory cortical neurons for task-relevant features, but the neurons show preserved and robust feature preference across learning and task-context. In contrast, neurons in the parietal cortex are recruited to increase selectivity for the rewarded features in a context-dependent manner. This combination supports both faithful representations of features in our environment and flexible adaptive action towards rewarding features.

## Methods

### Animals

All experimental procedures were carried out in accordance with institutional animal welfare guidelines and were licensed by the UK Home Office and are subject to restrictions and provisions contained in the Animals (Scientific Procedures) Act of 1986. Surgeries were performed on 20 adult C57Bl/6J wild-type (Charles River) and 7 PV-Cre mice (Jackson Laboratory, stock number: 008069; 20 male and 7 female) aged between 7-9 weeks. Mice were co-housed in a 12h reversed day/light cycle, and provided with standard diet, water and cage enrichment. During behavioral training and testing, animals were food restricted to maintain 85% or more of their free feeding body weights (2- 5 grams of food pellets/animal/day).

### Surgical Procedure

Mice were anaesthetized with 2-3% isoflurane (1-2% for maintenance) mixed with a 2 L/min oxygen supply and given pre-operative analgesia (Metacam, 5 mg/kg). Body temperature was closely monitored and maintained at 37 °C. Ointment was applied to the eyes (Xialin Night). After shaving the head, a circular piece of scalp was removed, and the underlying skull was exposed, cleaned, and dried. Animals were fitted with a headpost attached by dental cement (C&B or Universal Metabond) to allow head fixation during training and imaging. A single circular craniotomy (4 mm diameter) was created over V1 and PPC in the left hemisphere (stereotaxic coordinates from bregma: AP -4.1 mm, ML -2.5 mm for monocular V1 and AP -2.0mm, ML +1.7mm for PPC). After the exposure of the brain, V1 and PPC were located using their relative position from a reference point. A total of 1200 nL of virus (diluted 2:18) of GCaMP7s (Dana et al., 2019) (AAV-syn-jGCaMP7s-WPRE, Addgene: 104487-AAV1, in one animal GCaMP8f (Zhang et al., 2023), 162375-AAV9, AAV-syn-jGCaMP8m-WPRE) was injected in two sites in monocular V1 (ML -2.5 and ML -2.3) and a single site in PPC, 200 nL each at depths of 200 μm (layer 2/3) and 400 μm (layer 4) from the brain surface, using a pressure micro-injection system (Picospritzer III, Parker).The craniotomy was then sealed with a 4 mm-diameter glass coverslip. For some mice, a double-layer cranial window (two no. 1 thickness windows glued together with index-matched adhesive Norland #71) was used (Goldey et al., 2014). Animals were monitored for five days after the surgery with free access to food and water, while post-operative analgesia of (5 mg/kg) was provided on the first three days. After full recovery, mice were food-restricted and habituated to experimenter handling and head-fixation apparatus. Imaging and behavioral training started approximately three weeks after surgery to ensure stable viral expression.

### Behavioral Setup

Mice were head-fixed onto a Styrofoam cylinder (Figure 1A). Running speed on the cylinder was measured with an incremental encoder attached to the cylinder shaft (Kübler). A reward spout positioned near the snout of the mouse delivered a strawberry-flavored soy milk via a pinch valve (NResearch, UK), with licks being detected via a piezo disc sensor. Stimuli were presented on a 27” DELL U2715H monitor positioned at 45^°^ to the mouse body axis and 23 cm away from its right eye. Stimulus presentation and reward delivery were controlled with MATLAB using the Psychophysics Toolbox (Brainard, 1997).

Behavioral task trials were initiated by the mouse when running exceeded a threshold for a specified time (1-2 seconds). An intertrial interval (ITI), i.e. grey pre-stimulus illumination, was then presented for a randomized duration ranging between 3-25 seconds (3-6 seconds for testing) to discourage timed licking. To avoid excessive licking during training, the ITI was extended if mice licked during the grey screen stimulus, and 3-4 s timeout was given when mice licked during the unrewarded ‘no-go’ stimulus. Reward was delivered upon the first lick only during a reward zone (1-2 seconds after the rewarded ‘go’ stimulus onset), or through an auto-reward trigger at 2 seconds after stimulus onset if mice failed to lick. The stimuli consisted of static black and white gratings with a fixed spatial frequency (0.06 cpd) with 100% contrast and had either +45^°^ or -45^°^ orientation. Stimuli were presented in a randomized order and the rewarded stimulus was counterbalanced across mice.

### Experimental Protocol

Within 2-3 days, mice typically learned to run continuously on the cylinder and lick the spout for rewards without visual stimulation. During training, the animals learned to discriminate between the rewarded ‘go’ and non-rewarded ‘no-go’ stimulus (on average 3.5 weeks, range 1-8 weeks). Each training session was 60-90 minutes long and consisted of about 100-400 trials. Task performance was quantified for each session with behavioral d’ (Stanislaw and Todorov, 1999), with higher values reflecting better performance. Behavioral d’ was calculated by:

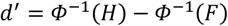

where *Φ* is the normal inverse cumulative distribution function, *H* is the rate of hit trials (licking for rewarded trials during the reward zone) and *F* is the rate of FA trials (licking for unrewarded trials during entire stimulus presentation). After the first or second training session, the pre-learning recording was performed. Once performance exceeded d’ > 2.0 for three consecutive training sessions, the post-learning recording was performed.

We used two-photon calcium imaging to measure the activity of the same layer 2/3 neuronal populations (150-200 *µ*m from the pial surface) in V1 and PPC while animals performed the visual discrimination task. ThorCam software (Thorlabs) was used to identify the injection site coordinates guided by blood vessel identification and relative locations of injection sites. Imaging was performed using a custom-built resonant scanning two-photon microscope and a Spectra Physics MaiTai eHP DeepSee laser (< 70 fs pulse width, 80 MHz repetition rate) at 920 nm using a Nikon 16x 0.8 NA objective. Images of 512 by 512 pixels and a field of view (FOV) of 500 × 500 µm were acquired using a 12 kHz resonant scanner (Cambridge Technology) and an FPGA module (PXIe-7965R FlexRIO, National Instruments), yielding an effective imaging frame rate of 30 Hz. Laser power was set between 16-25mW, and where possible, kept consistent throughout and across recording sessions. Retinotopic and orientation mapping helped identify a responsive FOV for recording.

Pre- and post-learning testing in the go/no-go discrimination task was similar to training (see above), except misses in go trials were not auto-rewarded and timeout was not given for FAs. Before each post-learning recording, we tried to identify the FOV from the pre-learning recording by matching anatomical landmarks. Mice, for which neither PPC or V1 was viable due to poor viral expression, overexpression, or bone regrowth under the window, were excluded from the study.

Immediately after the discrimination testing session, we typically determined the orientation tuning of all neurons in V1 and PPC, while mice ran freely on the cylinder without any trial initiation condition. We presented eight evenly spread static grating orientations, including the go and no-go stimuli, in randomized order (10 trials per stimulus). Grating stimuli were displayed for 1.5 seconds with 4-seconds-long ITIs.

## Data analysis

Image stacks were corrected for motion by maximizing the cross-correlation of all frames with a reference image (Guizar-Sicairos et al., 2008; Poort et al., 2015). Regions of interest (ROIs) were selected by manually assigning pixels to individual cells by inspecting individual frames, as well as the average and maximum projections of the imaging stacks. Pixels within each ROI were averaged to obtain raw fluorescence time series F(t). A causal moving average of 0.25 seconds was used to smooth F(t) and obtain baseline fluorescence F_0_(t) to determine the minimum value in the preceding 600 second time window of each time point. The difference between F and F_0_ was divided by F_0_ to compute the change in fluorescence relative to baseline, ΔF/F.

To analyze the responses of neurons to the grating stimuli, neuronal activity was aligned to stimulus onset, and a Wilcoxon signed-rank test was performed to determine if stimulus responses (the average ΔF/F in a time window of 1 second after onset) was significantly different from baseline activity, the average ΔF/F during 1 second preceding grating onset. Neurons for which this test was significant (p < 0.05) were classified stimulus responsive. To determine the stimulus preference of individual neurons, we computed a selectivity index (SI) from the mean responses to the go 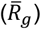 and no-go stimulus 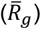 between 0-1 seconds, divided by the pooled standard deviation of the responses:

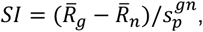

Where

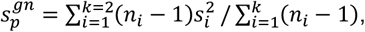

and *n*_*i*_ is the number of trials in condition *i* for *k* conditions. Thus, positive SI values indicate a bias in neural activity for the go stimulus and negative values a preference for the no-go stimulus. In Figure 2, we only included stimulus responsive neurons.

To match neurons across sessions, we automatically matched identical cells in the same FOV recorded on different days (Poort et al., 2015). This automatic matching was then validated manually for each cell. In Figure 2, since it was possible that some sites were recorded multiple times before or after learning, we averaged the selectivity index and mean PSTHs of matching cells both in pre-learning and post-learning sessions. When determining which cells were not modulated by behavior (in FA and CR trials in Figure 2F, or in Hit and Miss trials in Supplementary Figure 1B), we only included those cells that had similar responses in all sessions with trial number greater than 3 in each condition.

To test for significant differences between the selectivity indices before and after learning, a bootstrap test was used for partially independent data points (Efron, 1979). First, we calculated the number of cells included in the pre- and post-learning conditions. We then randomly selected the same number of cells in each condition, calculated the difference in the median of the selectivity indices of those cells pre-minus post-learning, and repeated the process 10,000 times. If the median was > 0, the p-value was the proportion of bootstraps below zero, and if the median was < 0, the p-value was the proportion of bootstraps above zero. We multiplied this p-value by two, as we conducted two-tailed tests.

For the analysis of matched cells (Figure 3), we calculated the correlation coefficient between the mean projection images of matching FOVs. We selected pre-post session pairs with maximum correlation coefficients, but only included those pairs for which this maximum was greater than 0.5. The Pearson correlation coefficient was calculated to quantify the correlations in Figure 3 and 4.

The preferred orientation was defined as the orientation with the maximum response (across eight orientations). A neuron was defined as tuned to a stimulus if the response to the preferred and orthogonal grating was significantly different (p < 0.05, Wilcoxon rank-sum test). We matched each orientation tuning block with the go/no-go task recorded on the same day. In some recording sessions, more than one orientation tuning block was conducted before or after the discrimination task, we only matched the first orientation block with the task. We calculated the (go/no-go) selectivity index in orientation tuning and then averaged it across sessions for matched cells in the same way as for the go/no-go task. However, since the p-value (and, hence, the preferred orientation) could not be averaged across multiple recordings, we determined the preferred orientation of the neurons from the session with the strongest tuning index (calculated as the difference between the preferred and orthogonal divided by the sum).

The proportions of switching neurons among ‘near go’ and ‘near no-go’ cells before and after learning (Figure 4C) were compared using bootstrapping. We compared all the groups (‘near go’ pre, ‘near go’ post, ‘near no-go’ pre, and ‘near no-go’ post) with one another in a pairwise manner. We selected two groups, and calculated the number of cells in each. Then, we randomly picked the same number of cells from the groups, calculated the proportion of switching neurons in each, and then subtracted one proportion from the other. We repeated the process 10,000 times. We conducted the same bootstrapping for all the pairwise comparisons (six comparisons in total). We calculated the p-values the same way as above, then corrected them using the Bonferroni method (i.e. multiplied each with the number of total comparisons). The bootstrap standard error of a group presented in Figure 4C was calculated from the standard deviation of the switching cell proportions from all the bootstraps and comparisons (thus, each group had 3 × 10,000 = 30,000 proportions in total).

To determine the effects of stimulus presentation and behavior on neuronal activity, we constructed generalized linear models (GLMs) in which the z-scored neuronal response (*R*) of each cell was regressed against the time series of different task events (Pho et al., 2018; Poort et al., 2015):

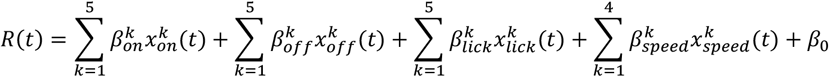

The calcium signal at frame *t* was modelled as the linear combination of the stimulus onset (*x*_*on*_), offset (*x*_*off*_), licking (*x*_*lick*_) and running speed (*x*_*speed*_) at the same frame. The stimulus onset was defined as a z-scored binary predictor convoluted with different decaying exponentials (timescale in frames *τ* = {5, 10, 20, 60, 200}). Thus, 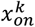 refers to the onset predictor convoluted with the *k*^*th*^exponential basis function (Poort et al., 2015). Similarly, stimulus offset and licking were provided to the model as z-scored binary predictors smoothed with the same five decay functions. Running speed was also z-scored and then convoluted with the following decay timescales specified in frames: *τ* = {4, 8, 20, 50}. Therefore, the full model contained 20 coefficients, which were estimated using ridge regression. The prediction accuracy was calculated by cross-validating the model with a test data including a random selection of 20% of the trials. We split the data 100 times, trained the model on the training data and calculated the model-explained variance following procedures similar to previous studies (Poort et al., 2015). The final variance (R^2^) was the average model performance across the 100 iterations. The stimulus-, licking- and running-explained variance was defined as the difference between the full model and a model lacking the stimulus, licking, or speed predictor, respectively.

## Acknowledgements

We thank all members of the Poort lab and the Beltramo lab for discussions. We thank Natsumi Homma for comments on the manuscript. This study was supported by a joint MRC DTP and Cambridge Trust studentship (M.V.), a UKRI Medical Research Council Equipment Grant (MC-PC-MR-X012271/1), and the Wellcome Trust (J.P., 211258/Z/18/Z).

## Author Contributions

J.M. and J.P. designed the experiments. J.M. and M.V. conducted the surgeries and collected the data. M.V. and J.M. analyzed the data with the help of J.P. All authors contributed to the writing.

## Supplementary Information

**Supplementary Figure 1.**
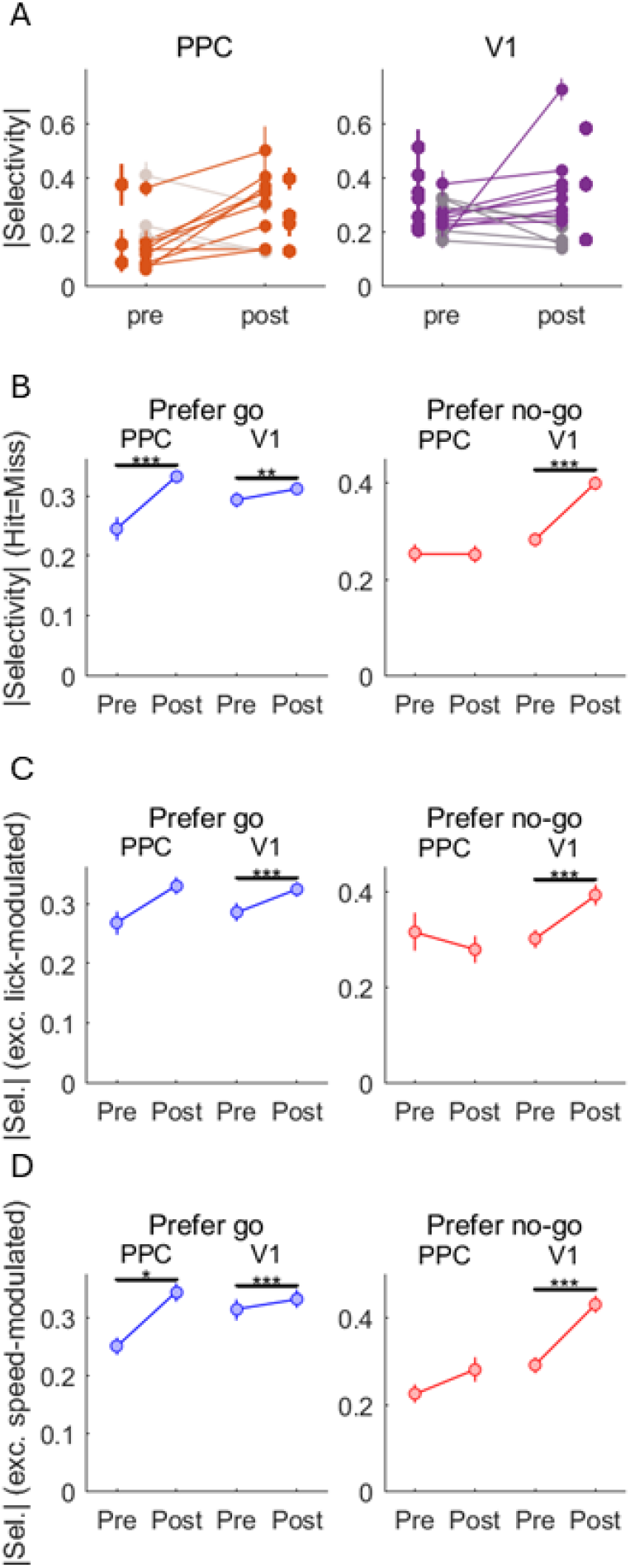
Selectivity increase is not attributed to variability across mice or motor-related effects. **A)** Average absolute selectivity across individual animals in PPC (left panel; only pre N=3, only post N=4, both pre and post N=12 mice) and V1 (right panel; only pre N=7, only post N=3, both pre and post N=13 mice). Solid lines, selectivity increased, faint lines, selectivity decreased across learning. Error bars, SEM. **B)** Similar to Fig. 2F, selectivity change of go- and no-go-preferring cells with equal Hit and Miss responses (PPC go-preferring cells, absolute selectivity pre-learning 0.25 ± 0.33, post-learning 0.33 ± 0.34, mean ± SD, N_pre_= 309, N_post_= 1007, p < 0.0001, bootstrap test; PPC no-go preferring cells absolute selectivity pre 0.25 ± 0.34, post 0.25 ± 0.42, N_pre_= 304, N_post_= 549, p = 0.44; V1 go-preferring cells, absolute selectivity pre-learning 0.29 ± 0.40, post-learning 0.31 ± 0.31, N_pre_= 1137, N_post_= 1073, p = 0.005; V1 no-go preferring cells absolute selectivity pre 0.28 ± 0.44, post 0.40 ± 0.46, N_pre_= 910, N_post_= 938, p < 0.0001). **C)** Selectivity change of go- and no-go-preferring cells after excluding the top 50% of neurons with greatest GLM R^2^ for licking. (PPC go-preferring cells, absolute selectivity pre-learning 0.27 ± 0.28, post-learning 0.33 ± 0.35, mean ± SD, N_pre_= 215, N_post_= 564, p = 0.08, bootstrap test; PPC no-go preferring cells absolute selectivity pre 0.32 ± 0.46, post 0.28 ± 0.48, N_pre_= 131, N_post_= 299, p = 0.32; V1 go-preferring cells, absolute selectivity pre-learning 0.29 ± 0.42, post-learning 0.32 ± 0.30, N_pre_= 722, N_post_= 599, p < 0.0001; V1 no-go preferring cells absolute selectivity pre 0.30 ± 0.46, post 0.39 ± 0.47, N_pre_= 574, N_post_= 466, p = 0.0002). **D)** Selectivity change of go- and no-go-preferring cells after excluding the top 50% of neurons with greatest GLM R^2^ for running. (PPC go-preferring cells, absolute selectivity pre-learning 0.25 ± 0.23, post-learning 0.34 ± 0.36, mean ± SD, N_pre_= 218, N_post_= 559, p = 0.008, bootstrap test; PPC no-go preferring cells absolute selectivity pre 0.23 ± 0.24, post 0.28 ± 0.48, N_pre_= 128, N_post_= 304, p = 0.20; V1 go-preferring cells, selectivity pre-learning 0.31 ± 0.47, post-learning 0.33 ± 0.35, N_pre_= 678, N_post_= 494, p = 0.0008; V1 no-go preferring cells absolute selectivity pre 0.29 ± 0.46, post 0.43 ± 0.46, N_pre_= 618, N_post_= 571, p < 0.0001). **B, C** and **D)** *p<0.05, **p<0.01, ***p<0.001, bootstrap test. Error bars, SEM.

**Supplementary Figure 2.**
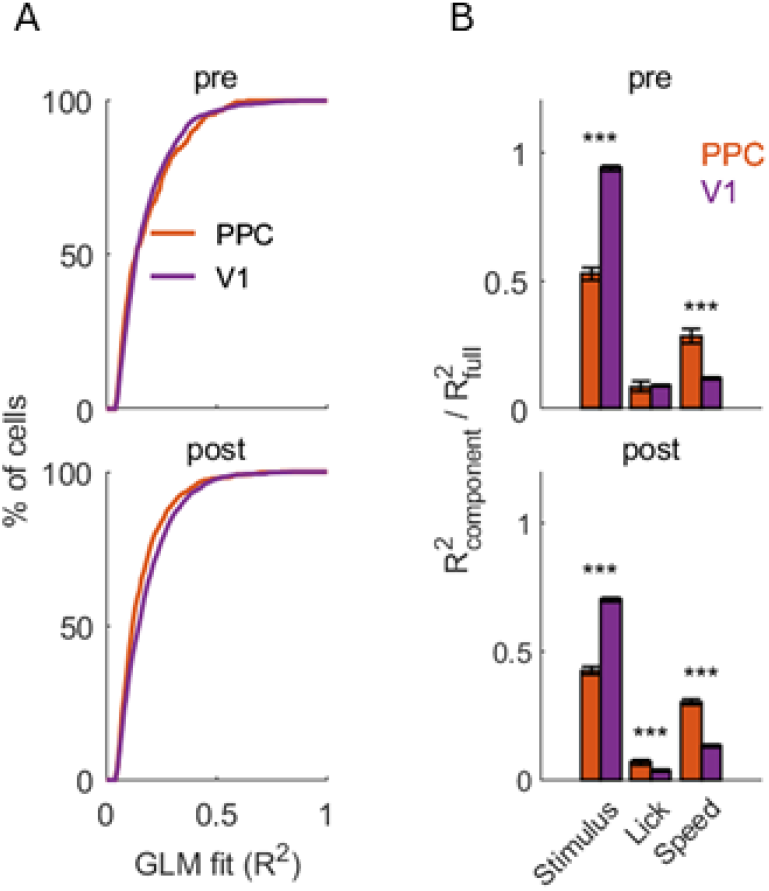
GLM analysis shows that V1 neurons primarily encode visual stimuli, while PPC neurons multiplex sensory and motor information. **A)** Cumulative histogram of the performance of the full GLM model in pre-learning (PPC 0.19 ± 0.13, mean ± SD, V1 0.18 ± 0.13, N_PPC_ = 273, N_V1_= 1824, p=0.74, Wilcoxon rank-sum test) and post-learning (PPC 0.16 ± 0.11, V1 0.18 ± 0.12, N_PPC_ = 648, N_V1_ = 1445, p=4.6 × 10^-6^). Only cells with R^2^ > 0.05 were included. **B)** R^2^ values for visual stimulus, licking and running relative to the full model in pre-learning (top panel; Stimulus: PPC 0.53 ± 0.41, V1 0.94 ± 0.46, mean ± SD, p=1.1 × 10^-47^, Wilcoxon rank-sum test; Lick: PPC 0.09 ± 0.34, V1 0.09 ± 0.16, p=0.43; Speed: PPC 0.28 ± 0.46, V1 0.12 ± 0.19, p=1.0 × 10^-11^) and post-learning (bottom panel; Stimulus: PPC 0.43 ± 0.40, V1 0.70 ± 0.32, p=1.9 × 10^-67^; Lick: 0.07 ± 0.26, V1 0.04 ± 0.15, p=9.4 × 10^-7^; Speed: 0.30 ± 0.28, V1 0.13 ± 0.21; p=8.2 × 10^-47^). Only cells with R^2^ > 0.05 for the full model were included. ***p < 0.001, error bars, SEM.

**Supplementary Figure 3.**
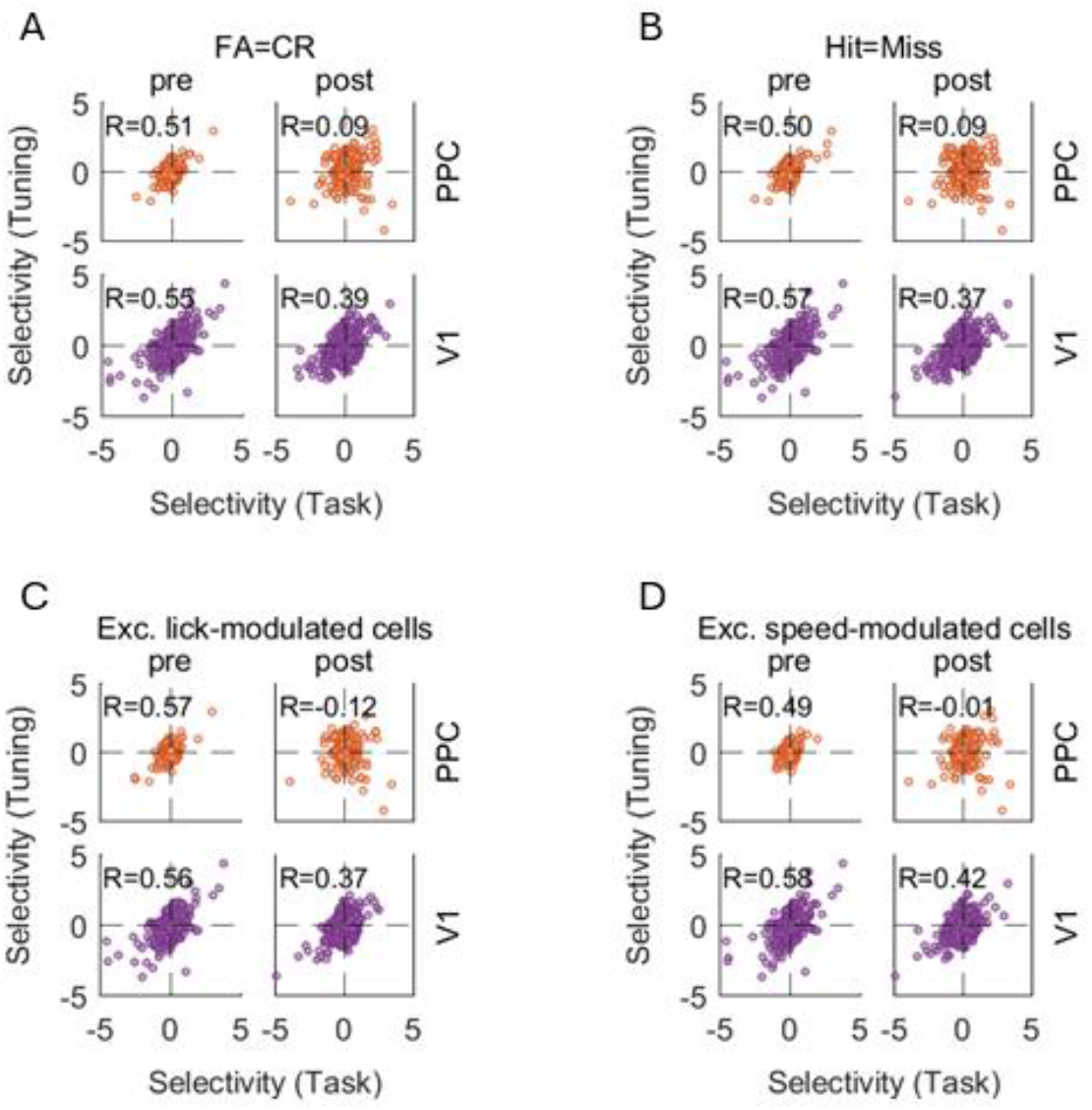
Context-dependent bias for the go stimulus in PPC after learning is not attributed to motor-related effects. **A)** Plot similar to Figure 4B, but with cells with equal FA and CR responses (PPC, pre: N=1070, p=4.6 × 10^-72^; PPC, post: N=2073, p=1.7 ×10^-5^; V1, pre: N=2180, p=4.8 × 10^-169^; V1, post: N=1976, p=2.8 × 10^-73^). **B)** Cells with equal Hit and Miss responses (PPC, pre: N=1161, p=4.1 × 10^-73^; PPC, post: N=2260, p=3.6 × 10^-5^; V1, pre: N=2107, p=4.2 × 10^-178^; V1, post: N=2343, p=2.4 × 10^-77^). **C)** Excluding top 50% cells with greatest GLM R^2^ for licking (PPC, pre: N=674, p=3.0 × 10^-60^; PPC, post: N=1245, p=1.1 × 10^-5^; V1, pre: N=1353, p=6.6 × 10^-115^; V1, post: N=1234, p=2.0 × 10^-40^). **D)** Excluding top 50% cells with greatest GLM R^2^ for running (PPC, pre: N=674, p=9.6 × 10^-43^; PPC, post: N=1245, p=0.629; V1, pre: N=1340, p=1.1 × 10^-120^; V1, post: N=1234, p=1.8 × 10^-54^).

## References

Andersen, R.A., Buneo, C.A., 2002. Intentional Maps in Posterior Parietal Cortex. Annu. Rev. Neurosci. 25, 189–220. 10.1146/annurev.neuro.25.112701.142922

Brainard, D.H., 1997. The Psychophysics Toolbox. Spat Vis 10, 433–436.

Colby, C.L., Goldberg, M.E., 1999. Space and attention in parietal cortex. Annu Rev Neurosci 22, 319–349. 10.1146/annurev.neuro.22.1.319

Dana, H., Sun, Y., Mohar, B., Hulse, B.K., Kerlin, A.M., Hasseman, J.P., Tsegaye, G., Tsang, A., Wong, A., Patel, R., Macklin, J.J., Chen, Y., Konnerth, A., Jayaraman, V., Looger, L.L., Schreiter, E.R., Svoboda, K., Kim, D.S., 2019. High-performance calcium sensors for imaging activity in neuronal populations and microcompartments. Nat Methods 16, 649–657. 10.1038/s41592-019-0435-6

Efron, B., 1979. Bootstrap Methods: Another Look at the Jackknife. Ann. Statist. 7. 10.1214/aos/1176344552

Felleman, D.J., Essen, D.C.V., 1991. Distributed Hierarchical Processing in the Primate Cerebral Cortex. Cereb. Cortex 1, 1–47. 10.1093/cercor/1.1.1

Glickfeld, L.L., Histed, M.H., Maunsell, J.H.R., 2013. Mouse primary visual cortex is used to detect both orientation and contrast changes. J. Neurosci. 33, 19416–19422. 10.1523/JNEUROSCI.3560-13.2013

Glickfeld, L.L., Olsen, S.R., 2017. Higher-Order Areas of the Mouse Visual Cortex. Annu Rev Vis Sci 3, 251–273. 10.1146/annurev-vision-102016-061331

Goard, M.J., Pho, G.N., Woodson, J., Sur, M., 2016. Distinct roles of visual, parietal, and frontal motor cortices in memory-guided sensorimotor decisions. eLife Sciences 5, e13764. 10.7554/eLife.13764

Goltstein, P.M., Coffey, E.B.J., Roelfsema, P.R., Pennartz, C.M.A., 2013. In vivo two-photon Ca2+ imaging reveals selective reward effects on stimulus-specific assemblies in mouse visual cortex. J. Neurosci. 33, 11540–11555. 10.1523/JNEUROSCI.1341-12.2013

Guizar-Sicairos, M., Thurman, S.T., Fienup, J.R., 2008. Efficient subpixel image registration algorithms. Opt. Lett. 33, 156. 10.1364/OL.33.000156

Harvey, C.D., Coen, P., Tank, D.W., 2012. Choice-specific sequences in parietal cortex during a virtual-navigation decision task. Nature 484, 62–68. 10.1038/nature10918

Henschke, J.U., Dylda, E., Katsanevaki, D., Dupuy, N., Currie, S.P., Amvrosiadis, T., Pakan, J.M.P., Rochefort, N.L., 2020. Reward Association Enhances Stimulus-Specific Representations in Primary Visual Cortex. Current Biology 30, 1866-1880.e5. 10.1016/j.cub.2020.03.018

Hovde, K., Gianatti, M., Witter, M.P., Whitlock, J.R., 2019. Architecture and organization of mouse posterior parietal cortex relative to extrastriate areas. European Journal of Neuroscience 49, 1313–1329. 10.1111/ejn.14280

Hubel, D.H., Wiesel, T.N., 1962. Receptive fields, binocular interaction and functional architecture in the cat’s visual cortex. The Journal of Physiology 160, 106–154. 10.1113/jphysiol.1962.sp006837

Itti, L., Koch, C., 2001. Computational modelling of visual attention. Nat Rev Neurosci 2, 194–203. 10.1038/35058500

Jurjut, O., Georgieva, P., Busse, L., Katzner, S., 2017. Learning Enhances Sensory Processing in Mouse V1 before Improving Behavior. J. Neurosci. 37, 6460–6474. 10.1523/JNEUROSCI.3485-16.2017

Krumin, M., Lee, J.J., Harris, K.D., Carandini, M., 2018. Decision and navigation in mouse parietal cortex. eLife 7, e42583. 10.7554/eLife.42583

Kukovska, L., Poort, J., 2024. The impact of parvalbumin interneurons on visual discrimination depends on strength and timing of activation and task difficulty. 10.1101/2024.06.07.597911

Lee, J.J., Krumin, M., Harris, K.D., Carandini, M., 2022. Task specificity in mouse parietal cortex. Neuron 110, 2961-2969.e5. 10.1016/j.neuron.2022.07.017

Licata, A.M., Kaufman, M.T., Raposo, D., Ryan, M.B., Sheppard, J.P., Churchland, A.K., 2017. Posterior Parietal Cortex Guides Visual Decisions in Rats. J. Neurosci. 37, 4954–4966. 10.1523/JNEUROSCI.0105-17.2017

Lyamzin, D., Benucci, A., 2019. The mouse posterior parietal cortex: Anatomy and functions. Neuroscience Research, Circuits and neural dynamics underlying behavior 140, 14–22. 10.1016/j.neures.2018.10.008

McNaughton, B.L., Mizumori, S.J.Y., Barnes, C.A., Leonard, B.J., Marquis, M., Green, E.J., 1994. Cortical Representation of Motion during Unrestrained Spatial Navigation in the Rat. Cereb Cortex 4, 27–39. 10.1093/cercor/4.1.27

Morimoto, M.M., Uchishiba, E., Saleem, A.B., 2021. Organization of feedback projections to mouse primary visual cortex. iScience 24, 102450. 10.1016/j.isci.2021.102450

Niell, C.M., Stryker, M.P., 2010. Modulation of Visual Responses by Behavioral State in Mouse Visual Cortex. Neuron 65, 472–479. 10.1016/j.neuron.2010.01.033

Niell, C.M., Stryker, M.P., 2008. Highly selective receptive fields in mouse visual cortex. J.Neurosci. 28, 7520–7536. https://doi.org/28/30/7520 [pii]; 10.1523/JNEUROSCI.0623-08.2008 [doi]

Pakan, J.M., Lowe, S.C., Dylda, E., Keemink, S.W., Currie, S.P., Coutts, C.A., Rochefort, N.L., 2016. Behavioral-state modulation of inhibition is context-dependent and cell type specific in mouse visual cortex. eLife Sciences 5, e14985. 10.7554/eLife.14985

Pakan, J.M.P., Currie, S.P., Fischer, L., Rochefort, N.L., 2018. The Impact of Visual Cues, Reward, and Motor Feedback on the Representation of Behaviorally Relevant Spatial Locations in Primary Visual Cortex. Cell Reports 24, 2521–2528. 10.1016/j.celrep.2018.08.010

Pho, G.N., Goard, M.J., Woodson, J., Crawford, B., Sur, M., 2018. Task-dependent representations of stimulus and choice in mouse parietal cortex. Nature Communications 9, 2596. 10.1038/s41467-018-05012-y

Poort, J., Khan, A.G., Pachitariu, M., Nemri, A., Orsolic, I., Krupic, J., Bauza, M., Sahani, M., Keller, G.B., Mrsic-Flogel, T.D., Hofer, S.B., 2015. Learning enhances sensory and multiple nonsensory representations in primary visual cortex. Neuron 86, 1478–1490. 10.1016/j.neuron.2015.05.037

Roelfsema, P.R., 2006. Cortical algorithms for perceptual grouping. Annu.Rev.Neurosci. 29, 203–227.

Saleem, A.B., Ayaz, A., Jeffery, K.J., Harris, K.D., Carandini, M., 2013. Integration of visual motion and locomotion in mouse visual cortex. Nat. Neurosci. 16, 1864–1869. 10.1038/nn.3567

Seabrook, T.A., Burbridge, T.J., Crair, M.C., Huberman, A.D., 2017. Architecture, Function, and Assembly of the Mouse Visual System. Annu Rev Neurosci 40, 499–538. 10.1146/annurev-neuro-071714-033842

Sorrell, E., Wilson, D.E., Rule, M.E., Yang, H., Forni, F., Harvey, C.D., O’Leary, T., 2025. An optical brain-machine interface reveals a causal role of posterior parietal cortex in goal-directed navigation. Cell Reports 44, 115862. 10.1016/j.celrep.2025.115862

Stanislaw, H., Todorov, N., 1999. Calculation of signal detection theory measures. Behav.Res.Methods Instrum.Comput. 31, 137–149.

Wang, Q., Burkhalter, A., 2007. Area map of mouse visual cortex. J.Comp Neurol. 502, 339–357.

Whitlock, J.R., 2017. Posterior parietal cortex. Current Biology 27, R691–R695. 10.1016/j.cub.2017.06.007

Zhang, S., Xu, M., Chang, W.-C., Ma, C., Hoang Do, J.P., Jeong, D., Lei, T., Fan, J.L., Dan, Y., 2016. Organization of long-range inputs and outputs of frontal cortex for top-down control. Nature Neuroscience 19, 1733–1742. 10.1038/nn.4417

Zhang, Y., Rózsa, M., Liang, Y., Bushey, D., Wei, Z., Zheng, J., Reep, D., Broussard, G.J., Tsang, A., Tsegaye, G., Narayan, S., Obara, C.J., Lim, J.-X., Patel, R., Zhang, R., Ahrens, M.B., Turner, G.C., Wang, S.S.-H., Korff, W.L., Schreiter, E.R., Svoboda, K., Hasseman, J.P., Kolb, I., Looger, L.L., 2023. Fast and sensitive GCaMP calcium indicators for imaging neural populations. Nature 615, 884–891. 10.1038/s41586-023-05828-9

